# Cell Storage Conditions Impact Single-Cell Proteomic Landscapes

**DOI:** 10.1101/2024.07.23.604834

**Authors:** Bora Onat, Amanda Momenzadeh, Ali Haghani, Yuming Jiang, Yang Song, Sarah J. Parker, Jesse G. Meyer

## Abstract

Single-cell transcriptomics (SCT) has revolutionized our understanding of cellular heterogeneity, yet the emergence of single-cell proteomics (SCP) promises a more functional view of cellular dynamics. A challenge is that not all mass spectrometry facilities can perform SCP, and not all labs have access to cell sorting equipment required for SCP, which together motivate an interest in sending bulk cell samples through the mail for sorting and SCP analysis. Shipping requires cell storage, which has an unknown impact on SCP results. This study investigates the impact of cell storage conditions on the proteomic landscape at the single-cell level utilizing a Data-Independent Acquisition (DIA) coupled with Parallel Accumulation Serial Fragmentation (diaPASEF). Three storage conditions were compared in 293T cells: (1) 37°C (control), (2) 4°C overnight, and (3) - 80°C storage followed by liquid nitrogen preservation. Either cold or frozen storage induced significant alterations in cell diameter, elongation, and proteome composition. By elucidating how cell storage conditions alter cellular morphology and proteome profiles, this study contributes foundational technical information about SCP sample preparation and data quality.

## Introduction

SCT is a powerful tool for investigating cellular heterogeneity. SCT has enabled the generation of tissue and organ atlases, such as the atlas of mouse organs (*Tabula Muris*^1^), in addition to human embryos^2^, and human skin^3^. While SCT effectively captures cell types and their change with pertrubations, transcript quantification is known to be highly biased and error-prone^4,5^, and transcripts are known to poorly correlate with proteins, which are the functional and structural units of cell systems. In contrast with the commoditization of SCT, SCP techniques are still being refined to achieve increased speed and sensitivity.

In recent years, researchers have made significant strides in the development of fast and sensitive SCP techniques^6–10^. Notably, a method leveraging diaPASEF^11,12^ with the timsTOF-SCP has enabled the quantification of over 1,000 proteins within a 15-minute total analysis time for each individual cell^13^. SCP has facilitated the identification and quantification of proteins within single cells, mostly in laboratory-cultured cell lines. Notably, diaPASEF was employed in the analysis of single muscle fiber proteome, with an output of 744 unique proteins derived from 8,764 unique precursors^14^.

Changes to the proteomic landscape due to sample processing is a key consideration in SCP. Prior to normal bulk proteomics analysis, it is common practice to freeze cells for extended storage durations. However, in SCP, cells must be sorted for sample preparation, and the impact of such storage conditions on cellular metabolism and identified proteins remains uncertain. Due to this requirement for cell sorting before SCP, and the lack of specialized sorting equipment at all institutions, we wondered how cold storage of bulk cells may influence data produced by SCP workflows. While it is established that cellular storage conditions exert influence on the proteome^15,16^, the extent to which these conditions affect the proteome at the single-cell level remains unexplored.

In this study, we determine the effects of cell storage conditions on 293T single cell proteomes using a CellenONE sorting and dual trap single column diaPASEF timsTOF SCP workflow. To investigate the effects of storage conditions on cellular proteomes, we subjected cells to three distinct storage conditions prior to proteomic analysis: 37°C controls, 4°C overnight in 5% DMSO (hereafter referred to as “4°C” or “cold”), and -80°C overnight in 5% DMSO at followed by storage in liquid nitrogen (hereafter referred to as “-80°C” or “frozen”). Our findings reveal that storage conditions exert disparate effects on individual cells at the proteomic level, underscoring the importance of avoiding cell storage via cold shock to ensure a more accurate representation of the original proteome. Our results show that both storage conditions affect proteins important for translation and metabolism, with some protein changes specific to either storage condition.

## Experimental Section

### Materials

HEK293T cells were a gift from the Van Eyk lab, heat inactivated Fetal Bovine Serum (FBS), high-glucose Dulbecco’s Modified Eagle’s Medium (DMEM), Antibiotic-Antimycotic, sterile Phosphate Buffered Saline (PBS) were purchased from Gibco/Life Technologies Corporation (Carlsbad, CA, USA). T75 cell culture flasks, 10% DDM, LC-MS grade DMSO were purchased from Thermo Fisher Scientific (Waltham, Massachusetts, USA). Cellenion X1 instrument was used for cell sorting (Lyon, France). 384-well plates were purchased from Bio-Rad (Hercules, CA, USA). Ultimate 3000 (Thermo Fisher Scientific) and Bruker timsTOF SCP (Billerica, Massachusetts, USA) instruments were used for LC-MS analyses. TEAB, Acetonitrile with 0.1% formic acid, LC-MS grade water with 0.1% formic acid were purchased from Millipore Sigma (Burlington, Massachusetts, USA). Sequencing grade trypsin was purchased from Promega (Madison, Wisconsin, USA). Cartridge trapping columns with a 0.17 µL media bed (EXP2 from Optimize Technologies) packed with 10 µm diameter, 100 Å pore PLRP-S (Agilent Technologies, Santa Clara. CA. USA) beads, and a PepSep 15 cm x 75 µm analytical column packed with 1.9 µm C18 solid phase (Bruker, Billerica, Massachusetts, USA) were used as the LC columns.

### Cell culture

HEK293T cell lines of passage number 10-13 were cultured in high-glucose DMEM supplemented with 10% FBS, 100 units/mL of penicillin, 100 μg/mL of streptomycin, and 250 ng/mL of Gibco Amphotericin B. Cell were cultured on Nunc T75 flasks and harvested through trypsinization and separated into 3 groups.

### Storage conditions

Approximately 10^6^ HEK293T cells were promptly subjected to sorting using the Cellenone instrument. A second cohort of 10^6^ HEK293T cells was suspended in fetal bovine serum (FBS) containing 5% dimethyl sulfoxide (DMSO) and stored at 4°C, while a third cohort underwent the same treatment but was subsequently stored at -80°C. Following overnight storage at -80°C, the cells were transferred to a liquid nitrogen storage system.

Subsequently, the second and third cohorts of cells underwent two washes in phosphate-buffered saline (PBS) and were then resuspended in PBS for sorting using the Cellenone instrument. This procedure brought samples to room temperature through the addition of PBS stored at room temperature.

### Cell sorting

The cellenONE system was utilized to dispense 200 nL of lysis buffer into each well of a Bio-Rad 384-well PCR plate. The lysis buffer composition consisted of 50 mM TEAB buffer, 0.2% DDM, and 200 µg/mL sequencing-grade trypsin. Following this, cells were resuspended in phosphate-buffered saline (PBS) to achieve a final concentration of approximately 3 × 10^5 cells/mL. SYTOX Green was added to the cell suspension at a final concentration of 25 µM, and the mixture was incubated in the dark for 5 minutes. Subsequently, cells were loaded into a piezoelectric dispensing capillary within the cellenONE X1 system for single-cell isolation. Cell viability was assessed using the green fluorescent channel of the system, with cells exhibiting SYTOX Green uptake being excluded from isolation. The 384-well plate holder was maintained at a temperature of 10°C using a water chiller throughout the sorting process. Following cell sorting, the plates were promptly transferred to a -80°C freezer for storage. Before subsequent analysis, 200nL of 40 µg/mL trypsin in 50 mM TEAB buffer was added in the wells and the sealed plates were incubated overnight in an oven at 37°C.

### LC-MS analysis

diaPASEF method was employed as described before by Kreimer et al.^13^ Assessment of system performance over injections was monitored by HeLa quality control (QC).

The chromatographic separation employed 0.1% formic acid in water as mobile phase A and 0.1% formic acid in acetonitrile as mobile phase B. The gradient profile was as follows: initially, 9% B at a flow rate of 500 nL/min; linear increase to 22% B over 8 minutes; further linear increase to 37% B over 4.7 minutes; followed by an increase to 1000 nL/min flow rate and 98% B over 0.2 minutes; maintaining 98% B for 1 minute; subsequent drop to 9% B at 1000 nL/min over 0.1 minutes; holding at 9% B at 1000 nL/min for 0.9 minutes; and finally returning to a flow rate of 500 nL/min in 0.1 minutes (totaling 15 minutes). The loading pump delivered 0.1% formic acid in water at a rate of 20 μL/min for the initial 6 minutes during trapping column cleaning, followed by a reduction to 10 μL/min from 6.5 to 15 minutes to load and desalt the subsequent sample. The valves and trapping columns were maintained at 55°C in the Ultimate 3000 column oven compartment, while the analytical column was kept at 60°C using the Bruker “Toaster” oven.

The autosampler was programmed to commence data acquisition immediately and to inject acetonitrile with 0.1% formic acid into the 20 μL sample loop, which was then passed through the trapping column using the loading pump flow. Following a second acetonitrile flush, the loop and needle assembly were rinsed with 25 μL of 0.1% formic acid in water. Subsequently, the autosampler aspirated 20 μL of 2% acetonitrile in water with 0.1% formic acid, which was dispensed and aspirated into the sample well three times to resuspend the sample, after which the sample was injected, and the needle assembly rinsed.

The dual trap single column (DTSC) configuration was adapted for single-cell analysis using cartridge trapping columns with a 0.17 µL media bed (EXP2 from Optimize Technologies) packed with 10 µm diameter, 100 Å pore PLRP-S beads (Agilent, Santa Clara, CA, US) and a PepSep 15 cm x 75 µm analytical column packed with 1.9 µm C18 solid phase (Bruker, Billerica, MA, US). The analytical column was directly linked to a 20 μm ZDV emitter (Bruker) integrated into the Bruker captive source. The capillary voltage was adjusted to 1700 V, with the dry gas flow set at 3.0 L/min and maintained at a temperature of 200 °C. Data acquisition was performed using DIA-PASEF. Ion accumulation and trapped ion mobility ramp was set to 166 ms. DIA scans were conducted with 90 m/z windows covering the range of 300–1200 m/z and 0.6–1.43 1/K0. A cycle time of 0.86 s was achieved, involving one full MS1 scan followed by 4 trapped ion mobility ramps.

### Proteomic data analysis

All Bruker .dia data were analyzed using DIA-NN 1.8.1^17^, using an in-house HEK293T library developed using a sequence of DDA data collected from FAIMS CV steps. DIA-NN settings were as follows: precursor m/z range: 300-1800, fragment ion m/z range: 200-1800, mass accuracy 10.0, MS1 accuracy 20.0, protease: Trypsin/P, missed cleavages: 1, protein inference: genes, neural network classifier: double-pass mode, quantification strategy: robust LC (high precision), crossrun normalization: global, library generation: smart profiling, and speed and ram usage: optimum results. The MaxLFQ^18^ calculated protein intensities from DIA-NN were used for this study. Missing values were replaced with zero. Scanpy version 1.9.1^19^ was used to perform single cell analysis on the resultant 1806 proteins and 189 cells across the three storage conditions, 37°C, 4°C and -80°C. Proteins that were detected in less than 10 cells were removed and cells with at least 200 proteins were retained, resulting in 1,269 proteins and 149 cells. Total count normalization was applied to make the sum of all protein intensities in a cell equal 10,000, followed by natural logarithm transformation of all intensities, regressing out unwanted variation on total counts, scaling quantities across proteins to have zero mean and standard deviation (SD) of 1, and repeating this same scaling method across all cells. All analysis was performed in Python version 3.9.12. Data was visualized using Matplotlib version 3.5.2^20^ and Seaborn version 0.12.2^21^.

### Statistical analyses

Single cell elongation and circularity values were exported from the cellenONE software. The normality of the data distributions was assessed using the Shapiro-Wilk test (Shapiro parameter) from the Scipy.stats module. The Kruskal-Wallis H test (Kruskal parameter) was employed to determine the statistical significance of differences between groups.

The principal component analysis (PCA) representation of the data was used to compute the neighborhood graph of cells, which was then clustered using Leiden^22^ and then embedded in a two dimensional space using Uniform Manifold Approximation and Projection (UMAP)^23^. Default Scanpy parameters were used except for changing Leiden resolution to 0.1.

Differential protein expression was tested using t-tests between each pair of conditions with 37°C (4°C and 37°C; -80°C and 37°C) using only non-zero protein quantities. A p-value was calculated only when there were at least three non-zero protein quantities per group for a protein. Benjamini-Hochberg(B-H) false discovery rate correction was applied to calculated p-values due to multiple testing. Log2 fold changes (FC) were calculated by subtracting the mean logged protein intensity between each pair of conditions for each protein. Only differentially expressed proteins (DEPs), or those with a B-H corrected p-value less than 0.01 and absolute value of log2(FC) greater than 1, were carried forward to pathway enrichment analysis. Statistical analysis was performed using SciPy version 1.11.4^24^.

Pathway enrichment analysis was performed on DEPs between 37°C and 4°C and 37°C and -80°C using GSEApy version 1.1.2^25^ in Python. The following human gene sets were used: GO Biological Process 2023, GO Molecular Function 2023, KEGG 2021 Human, Molecular Signatures Database (MSigDB) Hallmark 2020, and Reactome 2022. Enriched terms with an adjusted p-value less than 0.05 were selected and the -log10(adjusted p-value) of the top five terms was visualized in bar plots. Quantities of DEPs involved in two pathway, oxidative phosphorylation and translation, were visualized^78^.

## Data availability

All raw mass spectrometry data has been added to MassIVE database: ftp://MSV000095162@massive.ucsd.edu Password: hgu89!hgb

The spectral library from DDA is available from Zenodo: https://zenodo.org/doi/10.5281/zenodo.12802608

## Code and analysis availability

JupyterLab notebooks can be found at https://github.com/xomicsdatascience/Single-Cell-Storage.

The full processed data and analysis workflow to generate the clustering can be found at https://pscs.xods.org/p/wEx5D

## Results and Discussion

To understand how storage conditions apply to SCP and the heterogeneity of cells, we stored the cells under two conditions in addition to control freshly harvested cells. The standard procedure for the long-term storage of mammalian cell lines is to store the cells in fetal bovine serum (FBS) or growth media, with 5-10% DMSO in liquid nitrogen. We also tested 4°C as wet ice may be a more tractable alternative to frozen shipping. After cell storage, the cellenONE was used to sort single cells into 384-well cell PCR plates containing standard SCP lysis buffer (**Figure 1A**). The cellenONE has a standard camera to measure cell diameter, elongation and circularity, while a fluorescent camera prevents sorting cells positive for SYTOX Green, which only stains dead or unhealthy cells.

**Figure 1.**
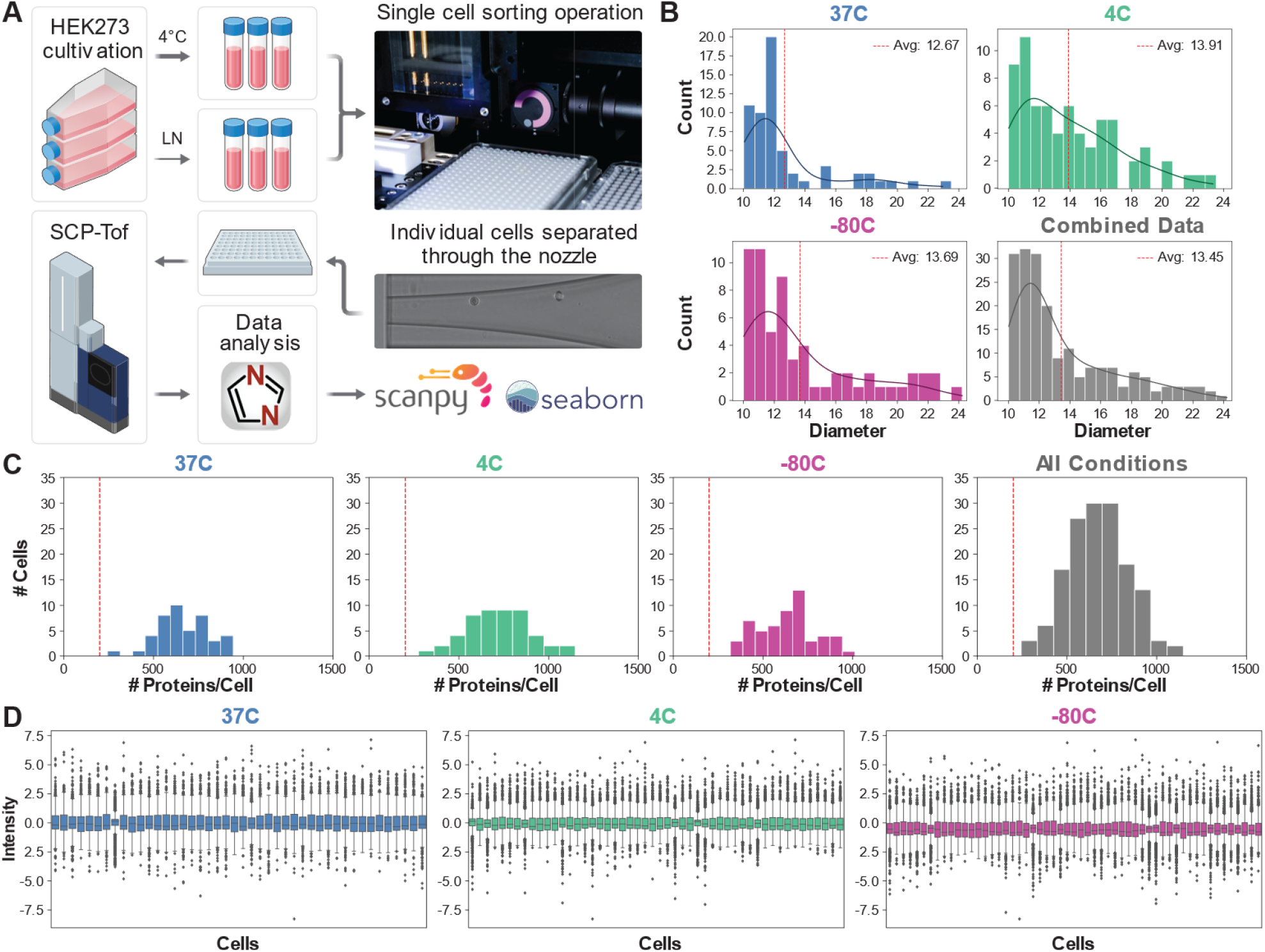
**(A)** Workflow overview diagram. **(B)** Histograms demonstrating cell diameters. Vertical dotted line denotes the average diameter while the solid blue line shows the KDE value. **(C)** Histograms of number of quantified proteins per cell in each of three conditions and when all cells are combined. Vertical dotted line indicates the minimum number of proteins per cell cut-off. **(D)** Boxplots demonstrating normalized protein intensity per cell analyzed across the three storage conditions. LN stands for ‘liquid nitrogen storage’.

We observed different cell population morphology across the three groups. The minimum cell diameter for selection was 10 μm, and the distribution of diameters was statistically different between groups (**Figure 1B**, Kruskal-Wallis Test p-value = 0.001). The cell elongation was also statistically different between the three groups **(Supplementary Figure 1**, Kruskal-Wallis Test p-value = 0.001), whereas the circularity was not different **(Supplementary Figure 2**, Kruskal-Wallis Test p-value = 0.1177). The maximum elongation and circularity value ranges were observed in the -80°C group, showing freezing conditions may influence cell shape more drastically. Cold temperatures are known to induce dehydration and cell shrinkage^26,27^. While sorting the cells, cells reach room temperature, thus the temperature of the surrounding PBS buffer may contribute to the cell size getting back to normal. DMSO also reduces the formation of water crystals, thus lowers the effects of dehydration.

The number of proteins quantified per cell was assessed in each of the three conditions as well when cells from all conditions were combined (**Figure 1C**). Only proteins quantified in at least 10 cells and cells with at least 200 proteins and were retained for downstream analysis. The mean number of proteins quantified per cell was significantly different among conditions by ANOVA (4°C: 722.6 proteins/cell, 37°C: 664.1 proteins/cell, -80°C: 633.4 proteins/cell; p=0.02). The distribution of proteins quantified per cell had a SD of 179.4 in the 4°C group, 160.3 in the -80°C group, and 145.5 in the 37°C group. After pre-processing (detailed in Methods), the distribution of all proteins per cell across technical batches are comparable with mean intensity per cell centered at zero (**Figure 1D**). Distribution of protein intensities per cell across conditions after only log2 transformation are shown in **Supplementary Figure 3**.

Using our 15 minute per cell dual trap-single column diaPASEF workflow^13^, we quantified 1,269 protein groups in 149 single cells across all experimental conditions. The clusters were colored by Leiden^22^, sample groups, and number of proteins quantified per cell after pre-processing in dimension reduced space by UMAP (**Fig 2A**). Plots of proteome profiles in dimension reduced space (UMAP) revealed that the Leiden clustering method^[19]^ suggests two groups, with the 37°C group separate from the other two groups. The Euclidean distances of the clusters in non-dimension reduced space were notably larger between the 37°C group and each the other two groups compared to -80°C vs 4°C (19.5 between 37°C and 4°C, 22.0 between 37°C and -80°C, and 13.7 between -80°C and 4°C). The protein counts for each group were randomly distributed within clusters. These findings suggest that storage conditions exert an influence on the proteomic composition of viable cells. In addition, we see a larger standard deviation of protein counts in the 4°C and -80°C groups, supporting that generally the storage conditions alter SCP data.

**Figure 2.**
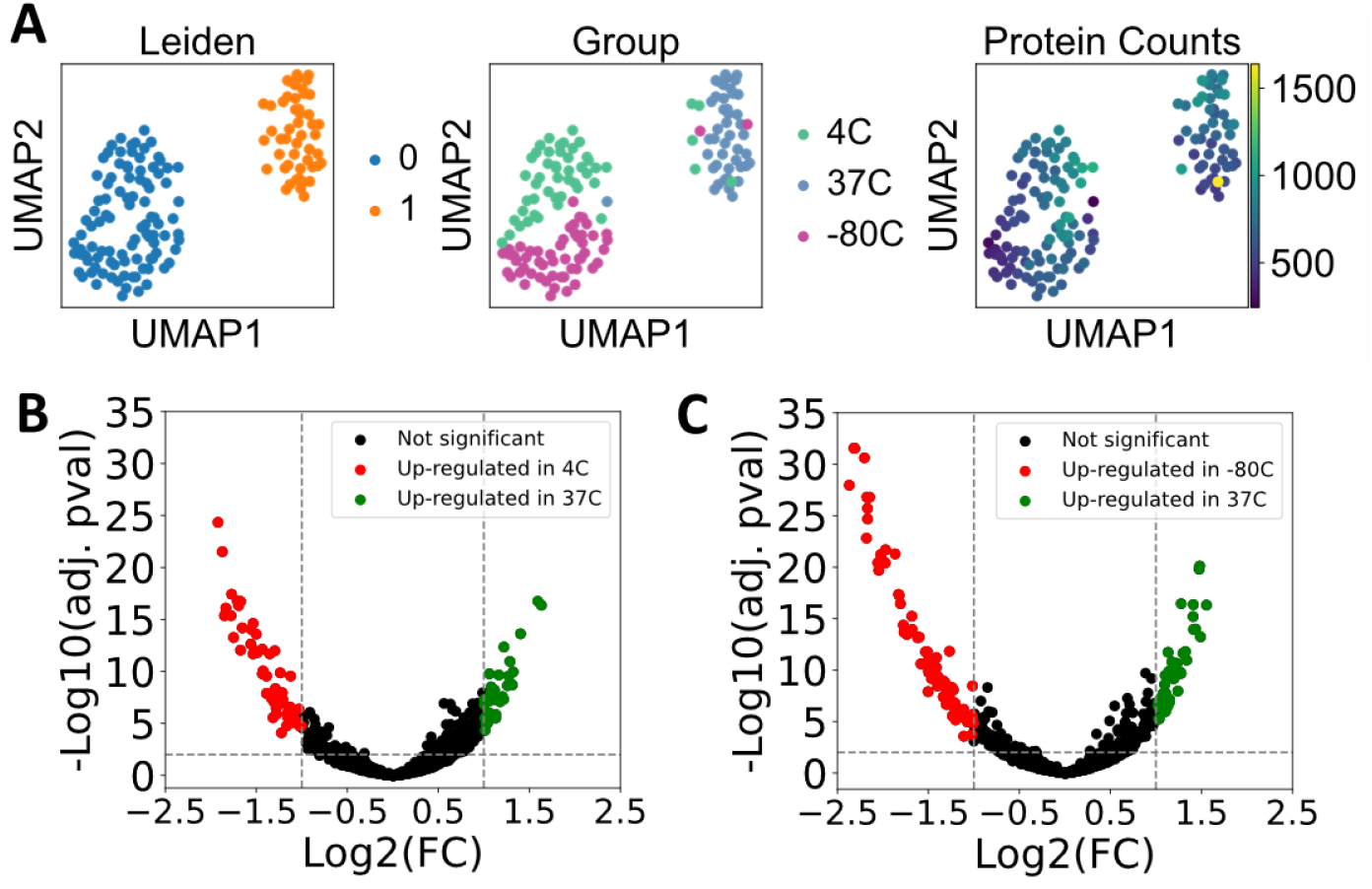
**(A)** Scatterplots of single cell proteomes reduced to 2D space with PCA followed by UMAP overlaid with Leiden clustering, storage condition group and number of proteins quantified per cell across the three experimental conditions. **(B)** Volcano plot depicting differential expression between the 4°C and 37°C groups, with Log2(FC) cutoffs set at -1 and 1, and -Log10 cutoff at 2. **(C)** Volcano plot illustrating differential expression between the -80°C and 37°C groups with the same cutoffs.

The proteome composition of each cold storage condition was compared statistically to the control using multiple t-tests and Benjamini-Hochberg correction by considering each single cell as a replicate measure. Volcano plots were used to visualize statistical differences and magnitude of change between 4°C and 37°C (**Figure 2B, Suppl. Table 1**) and -80°C and 37°C (**Figure 2C, Suppl. Table 2**), showing several proteins met cutoffs for adjusted p-value less than 0.01 and at least log2(FC) more than 1 or less than negative 1. In assessing the presence of technical bias that may cause quantities in one group to be consistently higher or lower than the other, we found the number of proteins with positive and negative log2(FC)s be balanced after using our data processing pipeline. We observed 64 upregulated and 33 downregulated proteins that met our cutoff described above in the 4°C group compared to the 37°C group. We observed 74 upregulated and 47 downregulated proteins in the -80°C group compared to the 37°C group.

### Enriched pathways summarize how storage conditions influence the proteome

The GSEApy interface to enrichr was used to determine statistically enriched pathways from: GO Biological Process, GO Molecular Function, KEGG, Molecular Signatures Database (MSigDB) Hallmark, and Reactome. The -log10(adjusted p-value) of the top five terms was visualized in bar plots for the total 97 proteins differentially expressed between 37°C and 4°C (**Figure 3A**) and 121 proteins differentially expressed between 37°C and -80°C (**Figure 3B**). Network maps were plotted to display the proteins associated with each pathway between 37°C and 4°C **(Figure 3C)** and 37°C and -80°C **(Figure 3D)**.

**Figure 3.**
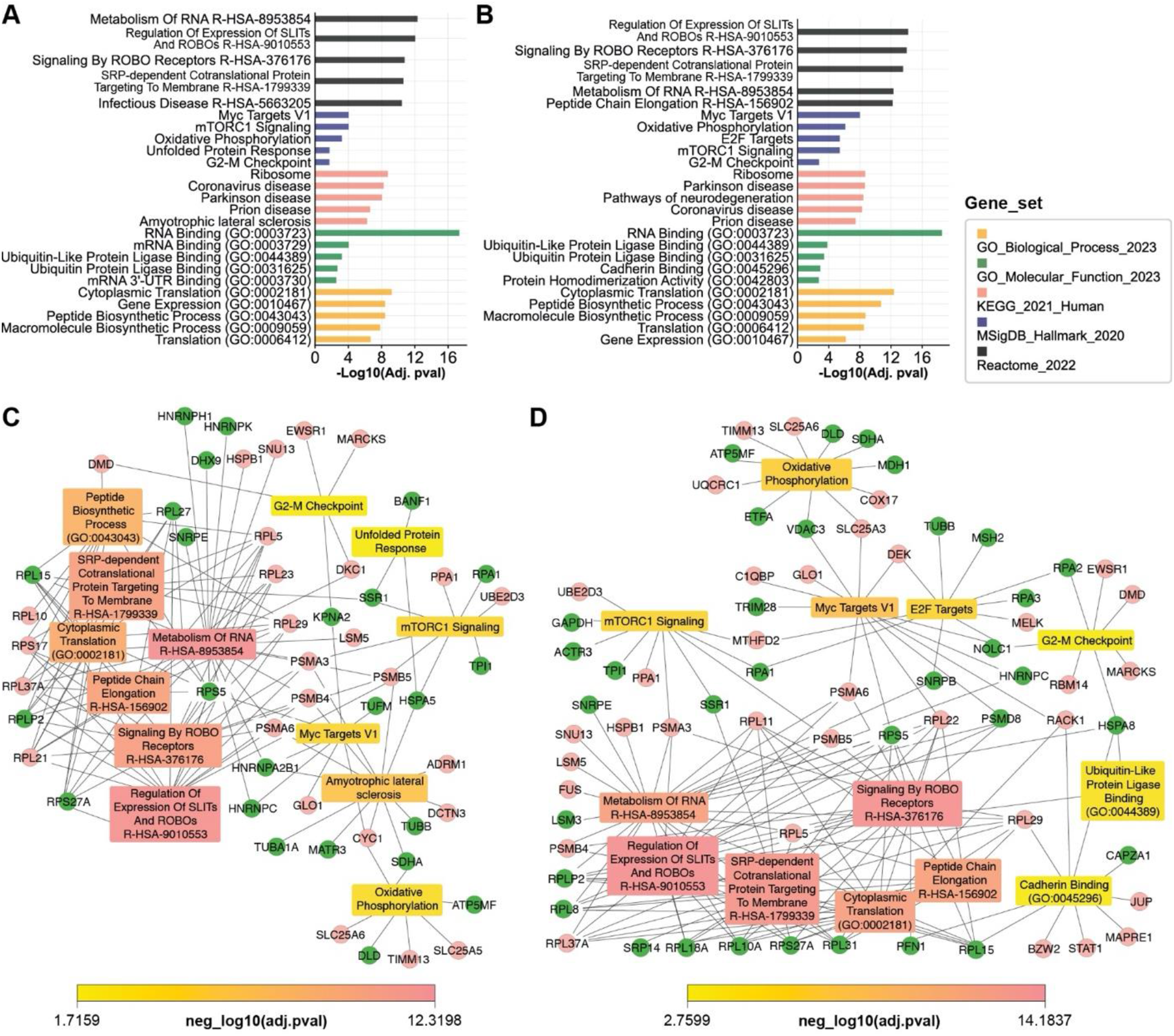
**(A)** Pathway mapping across 5 databases (GO Biological Process, GO Molecular Function, KEGG, MSigDB Hallmark, and Reactome) using differentially expressed proteins between 4°C group compared to the 37°C group. **(B)** Pathway mapping using differentially expressed proteins between - 80°C group compared to the 37°C group. **(C)** Cytoscape network map of differentially expressed proteins connected to enriched pathways between 4°C group compared to the 37°C group **(D)** and - 80°C group compared to the 37°C group. Green circle nodes display proteins that were upregulated in 37°C group, while pink were downregulated. The pathway boxes are colored according to the color scale for p-value significance. Only selected pathways that were discussed were included in network maps.

Differentially expressed proteins (DEPs) involved in each of the top 5 enriched pathways from each gene set are found in **Suppl. Table 3** (37°C vs 4°C) and **Suppl. Table 4** (37°C vs -80°C). Pathways enriched using only the proteins differentially upregulated in cells stored in 4°C compared to control are shown in **Suppl. Figure 4A** with DEPs significantly different between groups in each of the top 5 pathways per database listed in **Suppl. Table 5**; notable pathways included cellular responses to stress and cellular responses to stimuli. Pathways enriched using only the proteins differentially upregulated in cells stored in -80°C compared to control are shown in **Suppl. Figure 4B** with DEPs listed in **Suppl. Table 6**. This shows that the cells undergo stress due to DMSO or lower temperatures and this is reflected upon the proteome. Similar altered pathways were observed in the work of Verheijen *et al*. where the effects of DMSO on cellular processes was assessed.^28^ Even though <10% of DMSO in cryopreservation of human cell lines is considered to be non-toxic, studies showed that DMSO can alter crucial metabolic pathways, proven by proteomics, transcriptomics, and methylation analyses on human microtissues, and bacteria^29^. DMSO may be responsible for some proteomic alterations between the cold and frozen conditions relative to control cells.

Altered pathways include mTORC1 signaling, oxidative phosphorylation (OXPHOS), unfolded protein response, G2-M checkpoint, MYC targets V1, signaling by ROBO receptors, and translation. For both cold and freezing conditions, we have observed a decrease in the expression levels of PSMA3 and PSMB5 proteins, which are integral components of the 20S proteasome complex (**Figure 3C, 3D**)^30^. The observed downregulation of the proteasome complex suggests a potential disruption in protein degradation pathways, particularly affecting the clearance of misfolded proteins. This disruption is known to elicit endoplasmic reticulum (ER) stress, a cellular response associated with the induction of apoptosis^31,32^. ER stress is often followed by unfolded protein response (UPR), one of the pathways enriched in our proteome comparison of 4°C to 37°C. UPR maintains the protein folding homeostasis in the ER, maintaining the capacity of ER to correctly fold proteins^33^. mTORC1 pathway, ER stress and oxidative stress are three pathways that regulate each other, and may reflect hypoxia^34^. Furthermore, we observed downregulation in translational machinery proteins TARS2 and SSR1. SSR1 protein, part of the Signal Sequence Receptor (SSR) complex located in the ER membrane, plays a role in protein translocation, and can function as a down regulator of translational machinery^35^. Finally, we observed downregulation of ribosomal proteins (RPSs and RPLs)^36^.

The altered quantities of translation and OXPHOS proteins in either cold storage condition compared to control is shown in **Figure 4**. The mean of all non-zero protein quantities for each protein in the translation (**Figure 4A**) and OXPHOS (**Figure 4C**) pathways demonstrate the differential alteration of proteins with cold shock within the same pathway compared to control.

**Figure 4.**
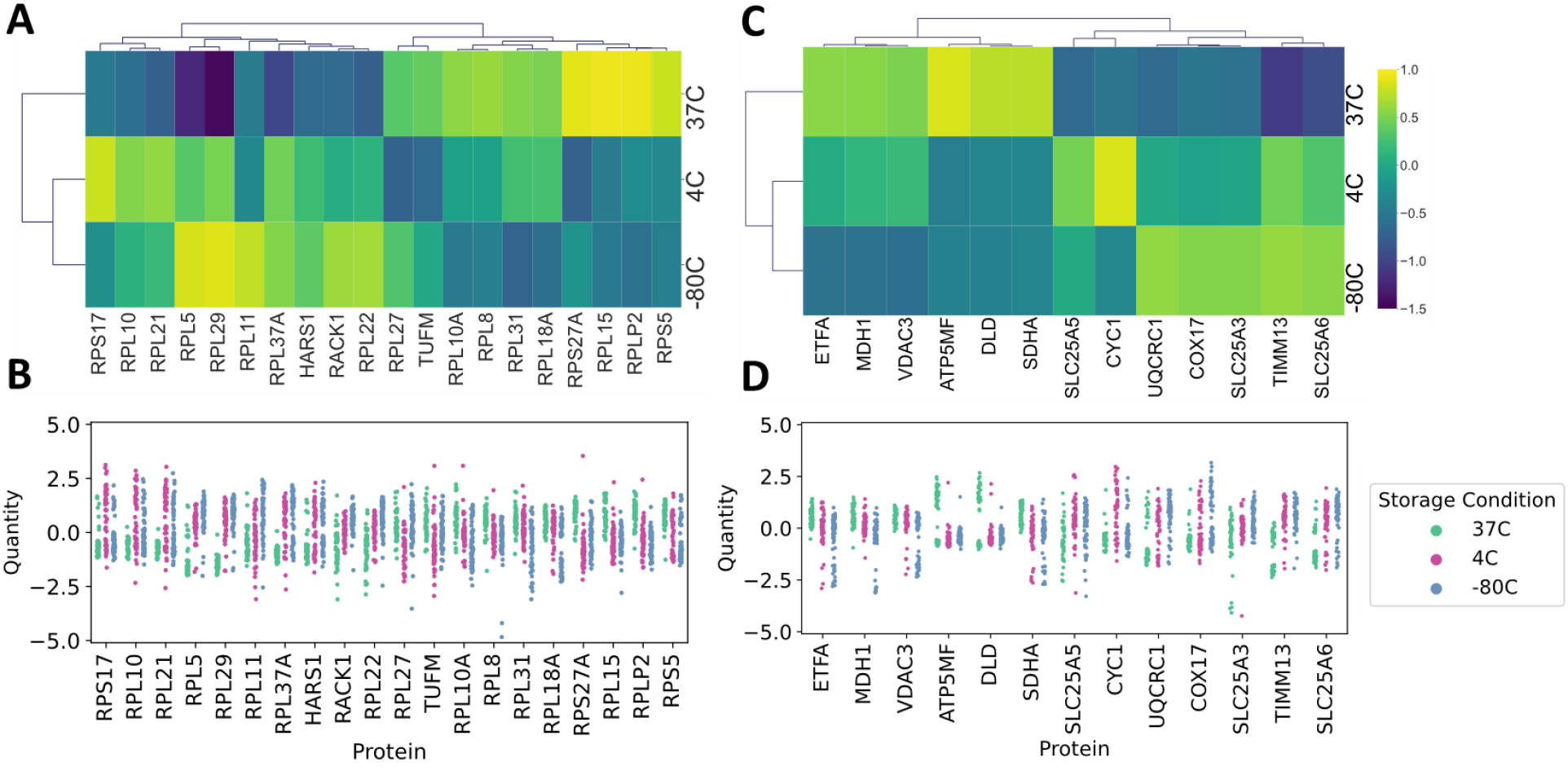
**(A)** Heatmap of average protein quantity for the DEPs in either cold condition vs 37°C involved in translation pathway. **(B)** Distribution of quantities from individual cells for DEPs involved in translation pathway across three conditions. **(C)** Heatmap of average quantity for DEPs in either cold condition vs 37°C involved in OXPHOS pathway. **(D)** Distribution of quantities from individual cells of DEPs involved in translation pathway across three conditions.

Interestingly, this differential alteration of proteins in the translation and OXPHOS pathways due to cold shock may influence other cellular processes, such as the G2-M checkpoint, which is a critical regulatory point where the cell assesses DNA integrity and determines whether to proceed with mitosis^37^. Even though our data do not display proteins that are directly involved in DNA integrity checkpoint, we observe an enrichment in this GO network in 4°C and -80°C groups, compared to the standard conditions. Even though the mechanism is not clear, short-term and long-term exposure of mammalian cells to hypothermia (4-10°C) are known to prevent cells from progressing beyond the G2-M checkpoint^38^, which could explain the enrichment in this GO pathway.

This suggests that the cellular response to cold shock may not only affect immediate protein pathways but also have broader implications on cell cycle regulation. Such interconnected influences highlight the complexity of cellular stress responses, particularly in relation to critical checkpoints like G2-M.

The proto-oncogene transcription factor MYC influences a wide array of genes that regulate various cellular functions, such as metabolism, proliferation, and morphology^39^. MYC interactome is known to comprise of at least 336 high-confidence proximity interactors, making this protein regulate a broad range of pathways in the cell^40^. MYC targets are linked to oxidative phosphorylation and the mTORC pathway in metabolic reprogramming^41^, suggesting that cold stress may modulate the metabolism of HEK293T cells. These insights into MYC’s role in regulating metabolic pathways further emphasize the multifaceted impact of cold stress on cellular function.

As we delve deeper into the specific pathways affected, such as the Slit/Robo pathway, it becomes evident how intricate and widespread the cellular response to cold stress can be. Our GO enrichment analysis revealed a significant enrichment in the Slit/Robo pathway, which plays a crucial role in regulating cell-cell and cell-matrix interactions^42^. It is plausible that cold stress alters these interactions, thereby inducing regulation of the Slit/Robo pathway. Given the pathway’s involvement in kidney morphogenesis, its enrichment in our study is understandable. Cells under cold stress are prone to apoptosis, and apoptotic cells may become hypercontractile, affecting cell-cell junctional stress, which could activate this pathway^43^. The results are highly heterogeneous, as displayed in the strip plot showing all log-transformed quantities of proteins (**Figure 4B, 4D**), indicating a diverse response among cells to the toxic effects of DMSO and hypoxic and cold stress, even though these cells were cultured immortal cells, showing lower diversity than cells isolated from tissues.

## Conclusion

In summary, we describe how cold or frozen cell storage before single cell sorting impacts SCP data. Our findings unveil notable alterations in cell diameter and elongation accompanied by significant proteome changes. Pathway enrichment analysis underscores cellular stress pathways induced by cold temperatures, with these effects persisting even during subsequent cell sorting procedures. Collectively, our results advocate for the preferential sorting of individual cells without subjecting them to prior storage conditions, particularly when aiming to preserve the native proteomic landscape of the cells for analysis. The main limitation of this study is that we cannot know from this data whether these changes are artifacts of the sample preparation or reflections of the true biological changes. Regardless, our data suggests that our view of single cell proteomes change with cell storage.

## Acknowledgements

This work was supported by NIGMS grant R35GM142502. Dasom Hwang helped with graphic design.

## ASSOCIATED CONTENT

### Supporting Information

**Supplementary Table 1**. Group means, Log2(FC), and -log10(adjusted p-values) for each protein between cells stored 4°C versus 37°C. Negative log2(FC) indicates mean expression was higher in 4°C group.

**Supplementary Table 2**. Group means, Log2(FC), and -log10(adjusted p-values) for each protein between cells stored -80°C versus 37°C. Negative log2(FC) indicates mean expression was higher in -80°C group.

**Supplementary Table 3**. Genes within each of the top 5 enriched pathways per database using all differentially expressed proteins between 4°C to 37°C.

**Supplementary Table 4**. Genes within each of the top 5 enriched pathways per database using all differentially expressed proteins between -80°C to 37°C.

**Supplementary Table 5**. Genes within each of the top 5 enriched pathways per database using all differentially upregulated proteins in 4°C compared to 37°C.

**Supplementary Table 6**. Genes within each of the top 5 enriched pathways per database using all differentially upregulated proteins in -80°C compared to 37°C.

**Supplementary Figure 1**. Distribution of intensities of all proteins per cell plotted across conditions when data is only log2 transformed.

**Supplementary Figure 2. (A)** Pathways enriched using only the proteins differentially upregulated in cells stored in 4°C compared to control. **(B)** Pathways enriched using only the proteins differentially upregulated in cells stored in -80°C compared to control.

## Author Contributions

According to CRediT taxonomy: **Conceptualization**, BO, SJP, JGM; **Methodology**, BO, AM, AH, JGM; **Software**, BO, AM, JGM; **Validation**, BO; **Formal Analysis**, BO, AM, JGM; **Investigation**, BO, AH, SY; **Resources**, BO, AM, YJ, JGM; **Data Curation**, BO, AM, JGM; **Writing – Original Draft**, BO, AM, JGM; **Writing – reviewing and editing**, BO, AM, JGM; **Visualization**, BO, AM, JGM; **Supervision**, JGM; **Project Administration**, JGM; **Funding Acquisition**, JGM.

